# A minimalist model lipid system mimicking the biophysical properties of *Escherichia coli’s* membrane

**DOI:** 10.1101/2024.09.29.615671

**Authors:** Nicolo Tormena, Teuta Pilizota, Kislon Voitchovsky

## Abstract

Biological membrane are highly complex systems that are of fundamental importance to the development and survival of organisms. Native membranes typically comprise different types of lipids, biomolecules and proteins assembled around a lipid bilayer structure. This complexity can render investigations challenging, with many studies relying on model membranes such as artificial vesicles and supported lipid bilayers (SLBs). The purpose of a model system is to capture the desired dominant features of the native context while remaining uniquely defined and simpler. Here, we search for a minimal lipid-only model system of the *Escherichia coli* inner membrane. We aim to retain the main lipidomic components in their native ratio while mimicking the membrane thermal and mechanical properties. We design a collection of candidate model systems reflecting the main aspects of the known native lipidomic composition and narrow down our selection based on the systems’ phase transition temperature. We further test our candidate model systems by independently measuring their elastic properties. We identify 3 ternary model systems able to form stable bilayers that closely mimic *E. coli*’s inner membrane lipid composition and mechanical properties. These model systems are made of commercially available synthetic 16:0-18:1 phosphatidylethanolamine (POPE), 16:0-18:1 phosphatidylglycerol (POPG), and 16:0-18:1 Cardiolipin (CL). We anticipate our results to be of interest for future studies making use of *E. coli* models, for example investigating membrane proteins’ function or macromolecule-membrane interactions.

**Significance Statement:** *Escherichia Coli* membranes serve as model systems for numerous fundamental and technological applications in the field of membrane biophysics. Being a Gram-negative bacterium, *E. Coli* is characterized by a thin cell wall that separates two phospholipid membranes: the inner and outer membranes. These membranes are complex, comprising many different lipids, proteins and other biomolecules. Here we develop a minimalist system to mimic the biophysical properties and lipidic composition of E. Coli’s inner membrane. Using only commercially available lipids, we develop a model membrane that can be used for studies where simplicity is needed to aid interpretation of the results, for example to investigate protein-induced mechano-transduction across E. Coli membranes.

## 1. Introduction

All types of organisms, from prokaryotic to eukaryotic, separate their internal environment from the exterior using biological membranes that consist of a self-assembled and self-synthesized double layer of phospholipids with a hydrophobic matrix in which a large number of proteins, and sugars, are bound or embedded (1). The function of biological membranes is multifold, first acting as a physical barrier, but also serving as a unique environment for certain types of biological processes such as the generation of electrochemical gradients of ions, one of the main energy sources of living cells (2), transport and motility (3). To do so, biological membranes have to sense, transmit, and respond to both chemical and mechanical stimuli. Chemical stimuli include the uptake and release of various molecules and ions (4) as well as sensing of the chemical composition outside the cell (5, 6). Mechanical stimuli arise from osmotic pressure changes (7, 8), changes in shear forces, cell-cell and cell-surface contacts (9, 10), as well as changes in the cytoskeleton (11, 12). A large number of macromolecules have been developed during cell evolution for the transduction of biomechanical stimuli, including proteins and glycolipids associated with the cell wall, the cytoskeleton, or the plasma membrane (13, 14). Proteins and lipids collaborate along this complex system to allow the transduction of any external stimuli and maintain cell physiological behavior.

The prokaryotic cell envelope consists not only of the cell membrane(s) but also of the above-mentioned cell wall, a structural layer made of a peptidoglycan matrix (15). In gram-positive bacteria, the cell envelope is composed of one plasma membrane and a thick external peptidoglycan layer, while gram-negative bacteria are characterized by a thinner cell wall that separates two phospholipid membranes called inner and outer membranes, respectively. One of the most common and well-studied gram-negative bacteria is *Escherichia coli*, which inner and outer membranes present a plethora of specific proteins relevant to physiological processes in each membrane. The outer membrane consists mainly of phospholipids and lipopolysaccharides (LPS) that act as endotoxins and play a fundamental role in the cell survival (16). It also contains tens of thousands of porin proteins allowing passive diffusion of molecules (17). In contrast, the inner membrane is composed of similar phospholipid families but with differing phosphate headgroup and acyl chain distribution (18), and with no LPS. It presents a lower phosphatidyl ethanolamine concentration and lower concentration of saturated lipid chains (18, 19). The inner membrane serves as a barrier for ions and consequently allows the generation of their electrochemical gradients. Some of the active membrane proteins are conserved in eukaryotic organisms (20), emphasising the importance of *E. coli*’s inner membrane as a model system. Over the last decade, a considerable amount of research has investigated *E. coli*’s membrane composition (21–24) and properties (18, 21, 22, 25–28), highlighting several important features. For example, the composition and biophysical properties are known to adapt to the environment (23, 24, 29–36). Additionally, *E. coli* membranes have compositional asymmetry, such as highly entropically disfavoured, unequal headgroup and acyl group asymmetries, both thought to be important for the biological function, but the origins of this asymmetry remain poorly understood (37). These features add to the challenge of developing a model *E. coli* membrane system since it may not be unique. Various studies have nonetheless attempted to do so, each with its own approach. Some of the simplest models omit cardiolipin, a key component of *E. coli* membrane (38, 39), while others use a wide range of phospholipid headgroups with different alkyl chain lengths (29–34), with or without cardiolipin. The biophysical properties of the model membranes are rarely investigated, which makes it difficult to assess whether the systems considered accurately reflect the intrinsic properties of *E. coli* membrane. For studies of biomolecules hosted in the model membranes this is problematic because phospholipids are crucial for the structural stabilization, active functionality, and localization of membrane proteins (40). Given the complexity of *E. coli* membranes and their dependence on the environment, even the best model system is unlikely to capture all of the native membrane’s properties, but model systems able to replicate the native lipidomic ratios and biophysical properties would already provide a valuable basis for a wide range of studies. The goal of such ‘minimal’ systems is to capture some key relevant properties of the natural system while remaining compositionally much simpler and well-defined (typically composed of artificial lipids and proteins mixed with precise stoichiometry (41)).

In this work, we systematically investigate binary and ternary lipid combinations of *E. coli*’s main lipids at stoichiometries close to those reported in lipidomic studies (18, 27, 42, 43). We aim to create a model membrane that replicates the *E. coli*’s inner membrane lipidic composition, thermal, as well as its mechanical properties at standard growth temperature (37°C). We combine commercially available synthetic lipids for their ease of use and widespread availability and characterize the properties of model systems by replicating the transition temperature (T_m_) (44) as well as comparing the in and out of plane stretching energies. Using a combination of differential scanning calorimetry (DSC), atomic force microscopy (AFM), and optical microscopy, we narrow down 18 possible combinations of the three main lipids present in *E. coli* and identify three mixtures that form stable and reproducible *E. coli* model membrane systems.

## 2. Materials and methods

All chemicals and lipids were obtained from commercial sources and used without further purification.

### Lipids

All the lipids were purchased from Avanti Polar Lipids (Alabaster, AL). The following lipids were purchased and dissolved in chloroform: 1-palmitoyl-2-oleoyl-sn-glycero-3-phosphoethanolamine (POPE), 1-palmitoyl-2-oleoyl-sn-glycero-3-phospho-(1’-rac-glycerol) (POPG), and 1’,3’-bis[1-palmitoyl-2-oleoyl-sn-glycero-3-phospho]-glycerol (CL). 1,2-dipalmitoyl-sn-glycero-3-phospho-(1’-rac-glycerol) (DPPG) was obtained in powder form. The native *E. coli* membranes (37°C growth) were obtained as *E. coli* Extract Polar (comprising only the polar lipids component) and *E. coli* Total Extract (full lipid extract) already dissolved in a chloroform: methanol solution.

### Chemicals

Salts (all >99% purity) were purchased from Sigma-Aldrich (Dorset, UK) and dissolved/diluted in ultrapure water (Merck-Millipore, Watford, UK). MOPS buffer-based solution was prepared with specific ion concentrations as follows: 50 mM NaCl, 9.5 mM NH_4_Cl, 0.5 mM MgCl_2_, 0.3 mM K_2_SO_4,_ and 1μM CaCl_2_-2H_2_O. The pH was adjusted to 6.5 prior to mixing with lipids.

### Large multilamellar vesicles preparation

Lipids dissolved in chloroform were mixed following the appropriate molar ratios into a 4 mL glass vial, -pre-dried under a gentle nitrogen flow, and fully dried overnight in a vacuum chamber. Large multi-lamellar vesicles were obtained by freeze-thawing (45, 46). Briefly, the lipid film was rehydrated in 2 mL of MOPS buffer-based solution to obtain a lipid concentration of 10 mg/mL and then briefly heated while sonicating in the sonication bath. Subsequently, the vial was frozen (left in the freezer for 15 min). This heating-freezing process was repeated for 6 consecutive cycles to successfully form large multilamellar vesicles (LMVs), which was confirmed by the lower turbidity of the solution and optical microscopy imaging.

### Unilamellar vesicles preparation

Lipids were mixed and dried following the same protocol for the LMVs preparation. The lipid film was subsequently rehydrated in 1mL of MOPS buffer-based solution obtaining a lipid concentration of 1mg/mL. The vial was gently bath sonicated for 15 min at a temperature 5-10°C higher than the highest T_m_ of the lipid species in the mixture, until the solution looked opaque and milky, indicating the formation of multilamellar vesicles. For small unilamellar vesicles (SUVs), the solution was extruded 31 times using a Mini-Extruder kit (Avanti Polar Lipids) with 1 Whatman 100 nm filter (GE Healthcare Life Sciences, Little Chalfont, UK).

### Supported lipid bilayers preparation

The SUVs solution was diluted 5 times to reach the 0.2 mg/mL concentration. 100 μL of SUVs solution were deposited on a disk of Grade 1 freshly cleaved Muscovite mica (SPI Supplies, West Chester, PA, USA) on the AFM stage and let incubating for 20 min at 50°C covered with a Petri dish. The sample was then gently rinsed with the MOPS-buffer based solution in order to remove any non-broken lipid SUVs and finally the temperature was cooled down to 40°C and equilibrated for 15 min, as a starting point for the measurement. This process ensures the formation of a spread and uniform supported lipid bilayers (SLB) system over the flat mica.

### Differential Scanning Calorimetry (DSC)

To observe the lipids main melting transition and extract the associated melting temperature values, DSC measurements were performed on a DSC 2500 (TA Instruments, Delaware, USA). Preliminary DSC heating tests were performed to identify ideal lipids concentration and DSC scan parameters, and to ensure satisfactory signal to noise ratio and reproducibility of the data (Fig. S1): with DSC, faster scan rates tend to provide a better signal to noise ratio. DSC test runs on binary lipid mixture were performed with increasing lipid concentration (from 1 mg/mL up to 10 mg/mL) and heating scan rate (from 2 °C/min up to 10 °C/min) while maintaining the temperature range of −10°C – 60°C (Fig. S1A, S1B). Since the scan rate can shift the experimental melting point, DSC cooling experiments were performed with the same increasing scan rates (from 2 °C/min up to 10 °C/min) while maintaining the same temperature range (Fig. S1C, S1D). This enables us to infer the melting point of our reference mixture at 0°C/min scan rate (thermodynamic equilibrium), and therefore estimate the effect of the scan rate on the experimental melting point (Fig. S1E). Practically, the dependence of the measured transition temperature on the scan rate is not trivial with previous studies (47, 48) reporting the following nonlinear dependence:

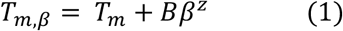

where *T*_*m,β*_ is the measured melting temperature at each scan rate, *T*_*m*_ is the equilibrium or ‘true’ melting temperature, *β* is the scan rate, and *B* and *z* are fitting parameters. Experiments run at different scan rates enabled us to determine *B* and *z*, and correct the measured transition temperatures (Fig. S1).

All other experiments were performed as follows: 10 μL of LMVs solutions at 10 mg/mL were loaded into the calorimeter and a heating rate of 5 °C/min was used in a temperature range of −10°C – 60°C. Samples were equilibrated for 5 min at the starting temperature (−10°C) before starting the measurement. 3 repeats were performed per each sample to enable statistical analysis.

### Optical brightfield microscopy

Images of LMVs were taken using Eclipse E200 (Nikon) microscope with 10x and 40x phase contrast objectives. Vesicles were imaged in a tunnel slide prepared as before (3, 49). Briefly, two parallel strips of double-sided sticky tape were positioned onto a microscope slide and covered with a 22×40 mm cover glass, which was pressed against the tape to form a tunnel. Approximately 10 µl of vesicles in solution was added to the tunnel for imaging. Pixel size was calibrated using a coverslip with 10 × 10 grid of 0.1mm squares (Graticules Optics).

### Atomic Force Microscopy

Imaging was conducted using a commercial Cypher ES AFM (Oxford Instruments, Santa Barbara, CA, USA), equipped with temperature control. SNL-10 cantilevers (Bruker Scientific Instruments, Billerica, MA, USA) with a nominal spring constant of 0.35 N/m were used. The tip has a pyramidal shape with a tip radius ≤ 12 nm at its apex. The AFM imaging was performed in amplitude mode, fully immersing the cantilever tip in the liquid. In this mode, the cantilever is acoustically oscillated at a frequency close to its resonance in liquid (∼10 kHz).

Force spectroscopy mapping was conducted in contact mode in liquid, using SNL-10 cantilevers. A schematics illustration of the measurement principle is shown in Fig. S4. A force map was created from 1024 force curves (32 × 32) over a 25 µm^2^ area. Calibration of the cantilever was performed by determining the inverse optical lever sensitivity by acquiring a force-distance curve on a stiff surface (mica) and the spring constant of each cantilever was determined from their thermal spectrum (50). This allowed for more accurate derivation of the Young’s modulus *Y* and membrane rupture force *F*_*r*_. Both *Y* and *F*_*r*_ were obtained using the same tip for all the measurements to ensure direct comparability between the results, regardless of any possible systematic offset. The emphasis is hence not placed on the absolute stiffness values (51, 52) but rather the relative differences between phases yielding the expected bimodal distribution. To minimize tip damage or contamination, AFM images were taken before and after the measurement, also ensuring similar membrane’s topographical features. The tip was cleaned with IPA and ultrapure water before starting a new measurement.

### Data analysis

DCS results were analyzed with the TRIOS Software, provided with the instrument. The software was used to correct thermogram baselines and then obtain *T*_*m*_ at the highest point of each calorimetric peak. DLS data were collected with the Zetasizer Family Software v.8.01, provided with the instrument. The size and of the vesicles and its uncertainty was obtained by fitting the experimental size histograms with a Gaussian distribution. AFM images and topographical AFM data were obtained and analyzed using the Gwyddion software (53), an open-source modular program for scanning probe microscopy data visualization and analysis. Optical microscopy images were analyzed using the open-source image processing package ImageJ/FIJI (54). Graphs were generated using Igor Pro Software (Wavemetrics, Lake Oswego, OR, US) and Python (55).

## 3. Results

### 3.1 PE, PG and CL are the main lipid species in *E. coli* membrane

The starting point for this study is a literature review of previous lipidomic studies of *E. coli* inner membrane to identify the main components and their relative fraction in the native membrane. This is consistent with our goal to create a model system that mimics the lipidomic composition of the native membrane. Further, to control the thermal and mechanical properties of the model system, the three most important parameters are: (i) the lipid polar head distribution, as it controls lipid-lipid interactions and charge distribution along the surface, (ii) the acyl chain length, as it controls membrane thickness, fluidity and membrane packing, and (iii) acyl chain saturation degree, because it regulates lipid packing within the bilayer. Taken together, these parameters constitute the major contributors to the specific mechanical properties of the bilayer and thus need to be correctly tuned.

The composition of *E. coli* ‘s inner membrane obtained from previous lipidomic and mass spectroscopy assays in summarized in Table 1, which also includes melting temperatures obtained from previous calorimetric studies (21, 22, 26). Because at *T*_*m*_ the thermal energy overcomes the internal energy of the membrane, *T*_*m*_ also offers a good indicator of how the membrane behaves mechanically: the internal energy of the membrane is the average inter-lipid interaction energy, influencing molecular order as well as dynamics within the membrane (44), and hence the propagation of any imposed mechanical stress.

**Table 1.**
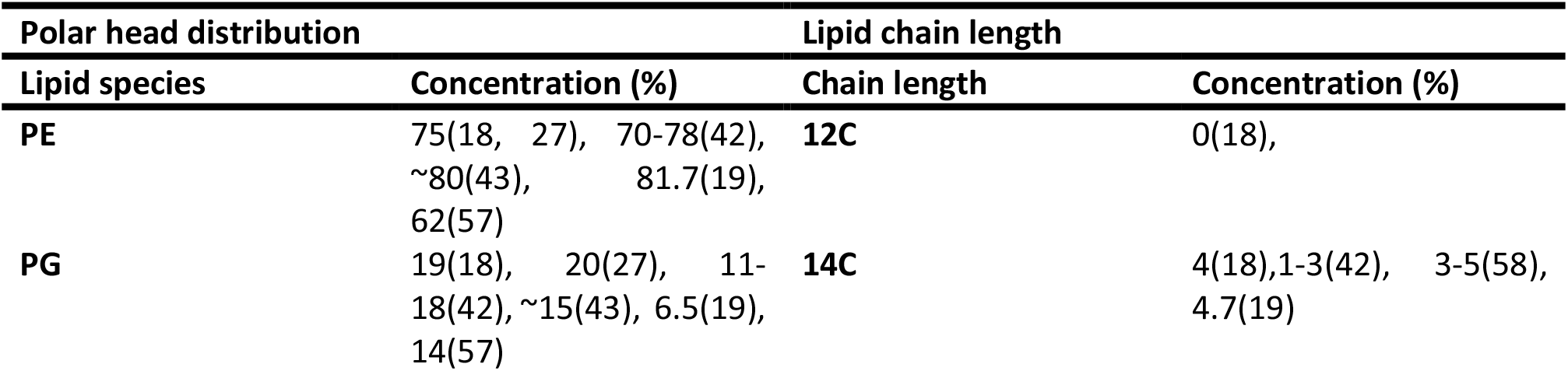
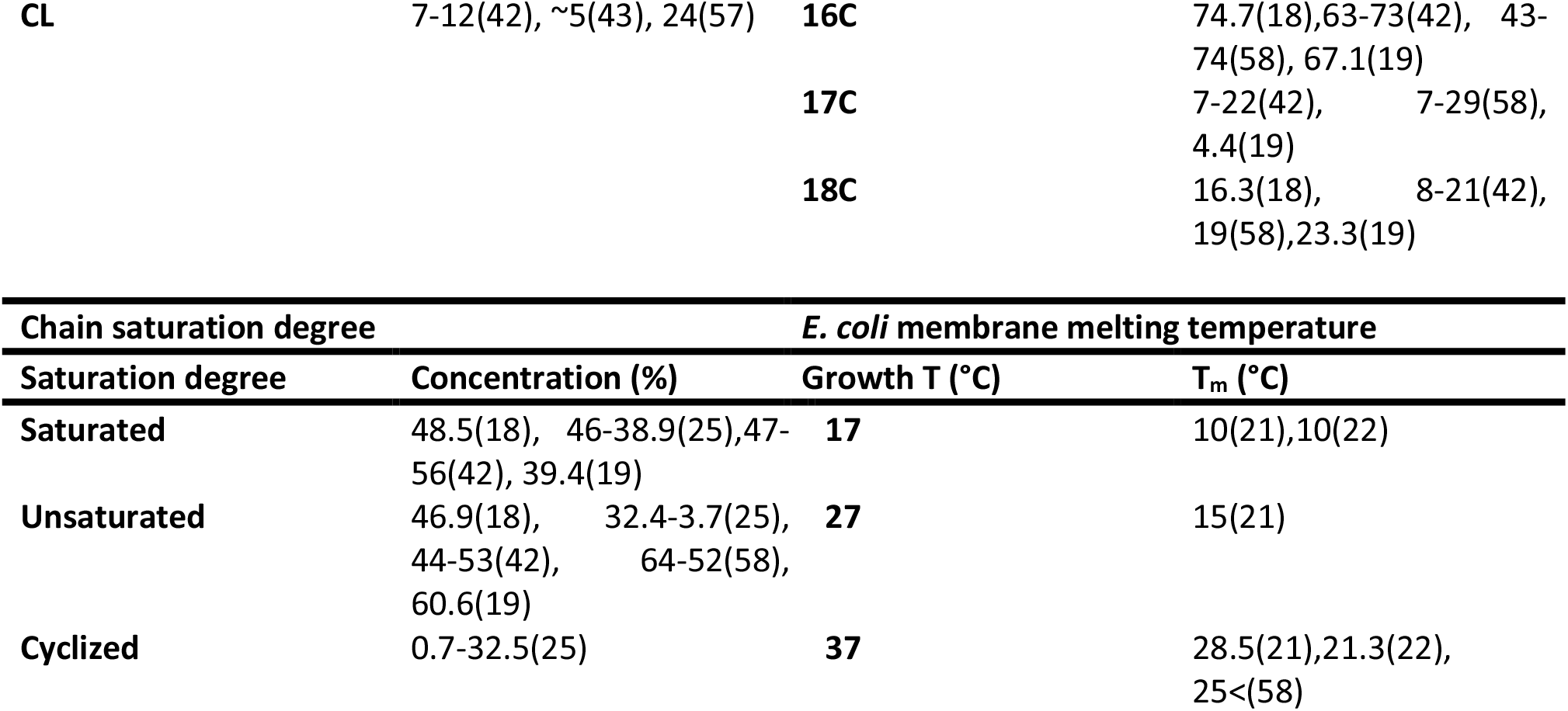
Summary of the E. coli inner membrane’s composition and properties, when grown at standard growth temperature of 37°C (upper part) and variation of E. coli membrane melting temperature based on growth temperature (lower part). The data compiles results obtained from published literature.

| Polar head distribution |  | Lipid chain length |  |
| --- | --- | --- | --- |
| Lipid species | Concentration (%) | Chain length | Concentration (%) |
| PE | 75(18, 27), 70-78(42), ~80(43), 81.7(19), 62(57) | 12C | 0(18), |
| PG | 19(18), 20(27), 11-18(42), ~15(43), 6.5(19), 14(57) | 14C | 4(18),1-3(42), 3-5(58), 4.7(19) |
| CL | 7-12(42), ~5(43), 24(57) | 16C | 74.7(18),63-73(42), 43-74(58), 67.1(19) |
|  |  | 17C | 7-22(42), 7-29(58), 4.4(19) |
|  |  | 18C | 16.3(18), 8-21(42), 19(58),23.3(19) |

**Table 1.** Summary of the *E. coli* inner membrane's composition and properties, when grown at standard growth temperature of 37°C (upper part) and variation of *E.
| Chain saturation degree |  | *E. coli* membrane melting temperature |  |
| --- | --- | --- | --- |
| Saturation degree | Concentration (%) | Growth T (°C) | T<sub>m</sub> (°C) |
| Saturated | 48.5(18), 46-38.9(25),47-56(42), 39.4(19) | 17 | 10(21),10(22) |
| Unsaturated | 46.9(18), 32.4-3.7(25), 44-53(42), 64-52(58), 60.6(19) | 27 | 15(21) |
| Cyclized | 0.7-32.5(25) | 37 | 28.5(21),21.3(22), 25<(58) |

The results obtained from previous calorimetric studies, which did not use any modification of the membrane such as the addition of markers (56), indicate a *T*_*m*_ value ranging from 7°C to 20°C lower than the growth temperature in the specific media. This depicts *E. coli* membrane as fluid and dynamic, rearranging the membrane composition through epigenetic reprogramming, e.g. in response to growth temperature, to shift its *T*_*m*_ and thus maintain overall fluidity across the range of conditions.

Here we focus on mimicking membranes at physiological growth temperature (37°C), meaning that the *T*_*m*_ of the model membrane should be lower than at least 30°C while maintaining the lipid polar headgroup ratios, chain’s length, and the overall degree of saturation as close as possible to that of the native membrane. From Table 1, the reported compositional ratios of *E. coli* inner membrane vary up to 20% (18, 27, 42, 43, 57). However, all results suggest that the main lipid species are phosphatidylethanolamines (PEs, 60-80% molar ratio) followed by phosphatidylglycerols (PGs, 15-30% molar ratio) and other minor lipids. Within these minor species, the most abundant is cardiolipin (CL, ∼5 %), which plays a crucial role in membrane’s physiological behaviour (59–61). CL has an unusual structure comprising 4 phosphatidyl chains connected through a glycerol linker (Fig. 1). The fact that a single headgroup is shared by 4 acyl chains confers unique physical properties to CL which can affect fluidity and charge density across the membrane (61). Previous studies have also reported a strong link between CL and membrane-protein interaction such as mechanosensitive channels (59). Furthermore, more than 700 protein-binding motifs have been identified along CL structure, which emphasizes the fundamental bio-functional role of this lipid in the plasma membrane despite its small concentration (60). We therefore include CL in our candidate model membranes. Apart from CL, most lipids in *E. coli* membrane show long carbon chains (>16C), with an even distribution between saturated and unsaturated lipid chains. Moreover, cyclized lipid chains are found in relatively high concentrations in the inner membrane (25), which significantly reduces bilayer density and lipid packing (29). Mass spectroscopy analysis have shown that the three most common lipid chains are C16:0, C18:1, and the cyclized C17:1 (cyC17:1) (28). Ideally, any *E. coli* model membrane should include these chains, but cyclized lipids are rare in other organisms and not commercially available. Keeping in our goal of a simple model system, we focus on more common non-cyclized lipids.

**Figure 1.**
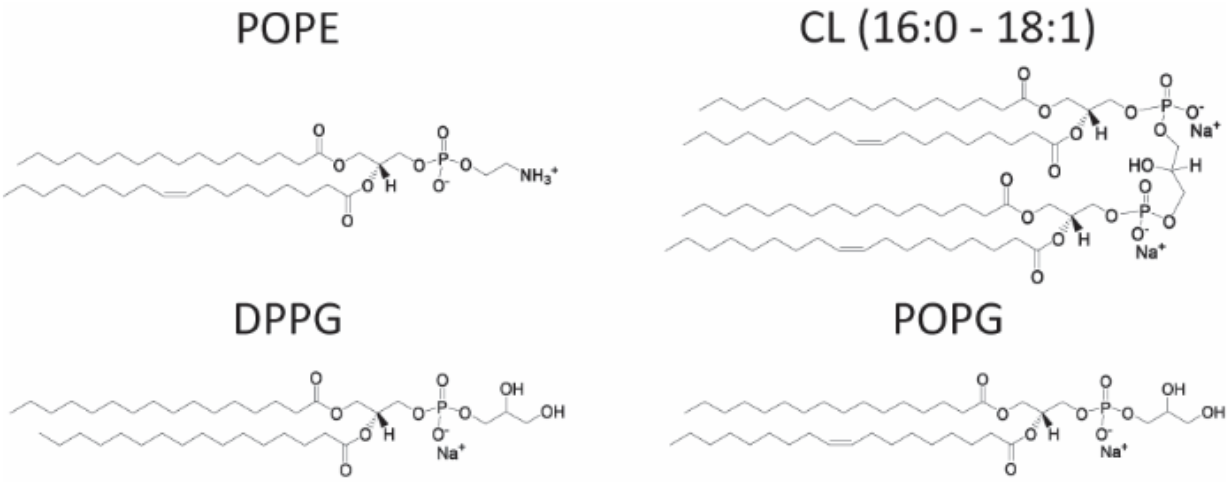
Schematic Lewis structure of lipids used to produce our lipid mixtures. Standard abbreviations have been used for the different lipid’s name. The molecular structures were reproduced from the information provided by Avanti Polar Lipid, the company from which the different lipids were purchased.

We identify 18 possible lipid mixtures that mimic the molar ratios of *E. coli* grown at 37 °C (Table 2). Specifically, we use binary and ternary mixtures in the appropriate molar ratios of POPE, POPG, DPPG and CL (16:0-18:1) because they all theoretically match the structural requirements of native *E. coli* membrane (Fig. 1). Both POPG and DPPG could work as the PG source for the *E. coli*-like model because they exhibit the most common acyl chains in these bacteria. POPG is often used for its relatively low *T*_*m*_ (−2°C), thus preventing phase separation in the membrane or the formation of ordered raft-like domains. In contrast DPPG has a *T*_*m*_ = 41 °C which is more likely to induce phase separation, but its two saturated acyl chains bring the overall molar ratio of unsaturated chains closer to the native ratio. It should be noted that the charged headgroup that characterizes the PG family could help prevent phase separation of DPPG, prompting us to keep both POPG and DPPG in our candidate model system.

**Table 2.** Candidate lipid mixture for the model systems for E. coli’s inner membrane when grown in physiological conditions (37 °C). Each mixture is of two or three types of phospholipids: POPE, POPG, DPPG and CL (16:0-18:1). Mixtures are indicated with a number and a letter, with the number indicating a specific lipid molar ratio and the letter indicating the specific PG lipid used (“A” being POPG and “B” being DPPG).

| Mixture name | POPE (%) | POPG (%) | DPPG (%) | CL (%) |
| --- | --- | --- | --- | --- |
| 1-A | 80 | 20 | 0 | 0 |
| 1-B | 80 | 0 | 20 | 0 |
| 2-A | 75 | 25 | 0 | 0 |
| 2-B | 75 | 0 | 25 | 0 |
| 3-A | 70 | 30 | 0 | 0 |
| 3-B | 70 | 0 | 30 | 0 |
| 4-A | 60 | 40 | 0 | 0 |
| 4-B | 60 | 0 | 40 | 0 |
| 5-A | 80 | 15 | 0 | 5 |
| 5-B | 80 | 0 | 15 | 5 |
| 6-A | 75 | 20 | 0 | 5 |
| 6-B | 75 | 0 | 20 | 5 |
| 7-A | 70 | 25 | 0 | 5 |
| 7-B | 70 | 0 | 25 | 5 |
| 8-A | 70 | 20 | 0 | 10 |
| 8-B | 70 | 0 | 20 | 10 |
| 9-A | 65 | 25 | 0 | 10 |
| 9-B | 65 | 0 | 25 | 10 |

### 3.2 POPG Ternary mixtures successfully mimic *E. coli* inner membrane transition temperature

Having identified candidate model membranes in Table 2, we next determine their melting temperatures as the first indicator of the average in-plane molecular interactions. For this purpose, we use lipidic LMVs samples in a MOPS buffer-based solution (see in Materials and Methods) and extract *T*_*m*_ for each sample from the main DSC transition peaks. The composition of MOPS matches the salt concentrations of commonly used *E. coli* growth media (62–64), but without the carbon source, making it a suitable environment for exploring model systems. And, LMVs are routinely used for this type of measurement because they enhance the signal-to-noise (SN) of the DSC compared to other types of lipid vesicles (45, 65–69). We selected a heating rate of 5 °C/min to ensure on optimal signal to noise ratio, and corrected for kinetic effects (47, 48) to infer the ‘true’ (i.e. thermodynamic equilibrium) *T*_*m*_ value (Materials and Methods, see also Fig. S1). Apart from each candidate mixture, we analyzed by DSC two *E. coli* inner membrane extracts as well, used here as references. Hereafter we refer to these references as *E. coli* ‘Native’ and *E. cli* ‘Polar Extract’. The first is a direct lipid extract of the *E. coli* inner membrane and the second is an extract further purified by removal of unknown lipid species from the native membrane, but still maintaining the complex mixture of PE and PG phospholipids with the original broad range of acyl chains. Both lipid mixtures were extracted from *E. coli* B (ATCC 11303) grown in Kornberg Minimal media at 37°C, as described by the commercial provider (see Materials Methods). We note that most studies on *E. coli* membrane lipidomic composition as well as majority of modern-day microbiology studies are on K-12 strain isolates. These two commonly used *E. coli* strains present highly similar genomes, which mainly diverge in their proteomic profiles (70, 71), but no alterations have been detected in their lipidomic profile nor on the key proteins involved in phospholipid synthesis, confirming the highly conserved composition of *E. coli* membrane within different strains. Table 3 summarises the DSC results of our mixtures with and without the previously described correction. Predicted *T*_*m*_ values calculated using weighted arithmetic mean (72) are also given where possible but only serve to identify the suitable temperature range for the measurements.

**Table 3.** Candidate lipid mixtures with melting temperatures theoretically calculated and experimentally evaluated with and without considering DSC scan rate effects. The reported experimental melting point temperatures are obtained through DSC experiments and the correction factor was calculated as described in Fig. S1E. Not all the theoretical T_m_ have been calculated since no information regarding the melting of pure CL is available to the best of our knowledge.

| Mixture name | Predicted $T_m$ (°C) | Experimental $T_m$ (°C) | Corrected $T_m$ (°C) |
| --- | --- | --- | --- |
| 1-A | 19.6 | $22.4 \pm 0.2$ | $20.7 \pm 0.2$ |
| 1-B | 28.2 | $30.8 \pm 0.1$ | $29.1 \pm 0.1$ |
| 2-A | 18.3 | $21.8 \pm 0.2$ | $20.0 \pm 0.2$ |
| 2-B | 29 | $30.3 \pm 0.2$ | $28.6 \pm 0.2$ |
| 3-A | 16.9 | $20.1 \pm 0.2$ | $18.4 \pm 0.2$ |
| 3-B | 29.8 | $31.7 \pm 0.2$ | $30.0 \pm 0.2$ |
| 4-A | 14.2 | $17.7 \pm 0.1$ | $16.0 \pm 0.1$ |
| 4-B | 31.4 | $34.7 \pm 0.4$ | $33.0 \pm 0.4$ |
| 5-A | - | $22.6 \pm 0.2$ | $20.9 \pm 0.2$ |
| 5-B | - | $29.0 \pm 0.3$ | $27.3 \pm 0.3$ |
| 6-A | - | $22.9 \pm 0.2$ | $21.2 \pm 0.2$ |
| 6-B | - | $30.7 \pm 0.2$ | $29.0 \pm 0.2$ |
| 7-A | - | $25.2 \pm 0.2$ | $23.5 \pm 0.2$ |
| 7-B | - | $31.2 \pm 0.2$ | $29.5 \pm 0.2$ |
| 8-A | - | $23.5 \pm 0.2$ | $21.8 \pm 0.2$ |
| 8-B | - | $30.3 \pm 0.1$ | $28.6 \pm 0.1$ |
| 9-A | - | $22.3 \pm 0.3$ | $20.6 \pm 0.3$ |
| 9-B | - | $31.2 \pm 0.2$ | $29.5 \pm 0.2$ |
| *E. coli* Native | - | $22.7 \pm 0.3$ | $21.0 \pm 0.3$ |
| *E. coli* P. Extract | - | $20.7 \pm 0.4$ | $19.0 \pm 0.4$ |

Fig. 2A-C visually summarises all our candidate samples and Fig. 2D shows the results from our DSC analysis for each of them. *E. coli* Native and *E. coli* Polar Extract mixtures exhibit *T*_*m*_ = 22.7 ± 0.3 °C and *T*_*m*_ = 20.7 ± 0.4 °C, respectively. The result is in line with previous lipidomic studies (Table 1) and used here as a guide to set the desired *T*_*m*_ we expect from our model system. Mixtures containing POPG shows a significantly lower melting point compared to those with DPPG with an average 8 to 10°C gap between equivalent mixtures. These differences are in line with the significant *T*_*m*_ difference between the two pure lipid species and suggest their homogeneous mixing in POPE. The presence of a unique main peak in the DSC curves (characterized by an average change in enthalpy of 20-40 kJ/mol), generally associated with the melting of a lipid membrane (73) (see Fig. S2), further confirms homogenous mixing. All mixtures exhibit a *T*_*m*_ lower than 37°C – our reference *E. coli* growth temperature – in principle making all the mixtures still eligible for our model. Ternary mixtures exhibit more complex behavior, without an obvious *T*_*m*_ trend emerging. This is in line with previous studies that investigated the phase transition of CL containing bilayers and found heterogeneous behavior. For example, CL molecules can elevate the *T*_*m*_ from −20°C in the absence of cations, up to 30°C in their presence – a *T*_*m*_ even higher than for their corresponding diacyl phosphatidylglycerols (74). Moreover, CL exhibits a small, poorly flexible structure that drives complex -and not yet fully understood-interactions with both different and similar lipids in the bilayer. Depending on the bilayer composition, CL can be homogenously dispersed within the bilayer (75), or separate in CL-enriched domains (76). The highly heterogeneous and not fully understood nature of CL renders its effect on membrane fluidity challenging to predict and explain compared to simpler phospholipids.

**Figure 2.**
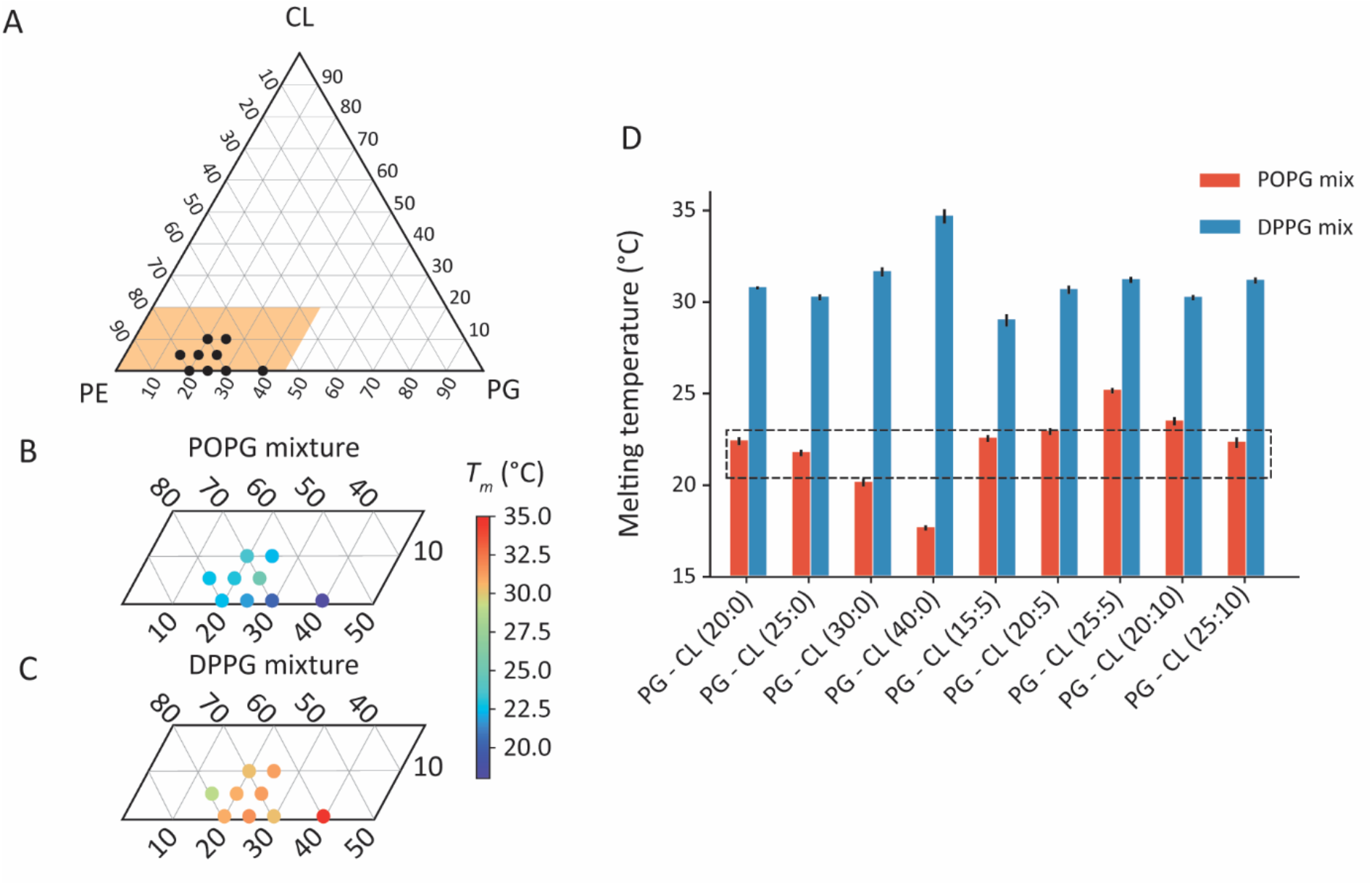
Melting temperature analysis of artificial lipid vesicles based on physiological E. coli inner membrane’s compositions. (A) Ternary diagrams of PE-PG-CL model systems explored showing the physiological composition range of E. coli inner membrane. (B-C) Zoom in the ternary diagrams of POPG (B) and DPPG (C) based mixtures together with the melting temperature (obtained as discussed in the text and Materials and Methods) in each case. (D) DSC results displaying differences between POPG and DPPG systems and highlighting the native E. coli inner membrane ranges based on literature and reference experiments (dashed box).

Comparing our results to both *E. coli* Native and *E. coli* Polar Extract mixture rules out the possibility of using DPPG as a PG source for our model membrane, presumably reflecting more ordered structures than in the native bilayers. However, mixtures with DPPG do not appear to phase separate despite their high *T*_*m*_ (single peaks in DSC, Fig. S2), but may form a molecularly well-packed homogenous membrane. The best match to our *E. coli* mixture references is obtained for 3 different ternary mixtures of phospholipids: 80% POPE, 15% POPG and 5% CL (composition #5A in Table2), 75% POPE, 20% POPG and 5% CL (composition #6A in Table2) and POPE 65%, 25% POPG and 10% CL (composition #9A in Table2).

It is worth noting that two binary mixtures match the native membrane behavior: 80% POPE and 20% POPG (composition #1A in Table2) and 75% POPE and 25% POPG (composition #2A in Table2). These mixtures still reflect molecular ratios of the most abundant lipids reported for *E. coli*, are able to reproduce the *T*_*m*_ of our reference samples, and might hence be sufficient for some studies provided the other biophysical properties also match those of the native membrane. However, this would need some caution since the lack of CL that could significantly influence any protein-related studies (59). It is also worth noting that the most recent study on the PE composition of *E. coli* membrane, reports about 60% PE for the inner membrane, and claims previous estimates of 75-80% are an overestimate (57). In this scenario, and particularly if further confirmed by future studies, our results would suggest a unique candidate for *E. coli* model membrane, which is POPE 65%, 25% POPG, and 10% CL (composition #9-A in Table 2). We emphasize that the model systems do not take into account the role of proteins and other biomolecules lodged into the *E. coli* membranes but rather offer a lipid matrix reflecting the native lipid composition while simultaneously retaining some key biophysical properties.

### 3.1. Mechanical properties of the candidate and *E. coli* membranes

Before measuring the mechanical properties of our candidate model systems, it is necessary to demonstrate that the mixtures can indeed form stable and homogenous unilamellar vesicles, as well as smooth, stable supported lipid bilayers. The formation of stable and homogenous SLBs from SUV deposition is not trivial because CL can affect the bilayer fluidity, evolution and membrane packing (77), sometimes inducing some molecular rearrangement over time and precluding the formation of stable flat SLBs (78). Additionally, the formation of SLBs with PE and PG has been previously reported as challenging due to the negative charge of the PG headgroups, the conformation of POPE/POPG molecules (79) and the effect of PE lipids on membrane curvature (80). To confirm the formation of stable and homogenous SUVs we used optical microscopy, illustrated in Fig. 3A-B for LMVs of composition #6-A (Table 2), revealing rounded vesicles which were stable for at least 14 days. Similarly, AFM imaging of supported candidate membranes (Fig. 3C) reveal smooth, stable patches. Here, SLBs were formed on an atomically flat mica using extruded 100 nm SUVs (see *Materials and Methods*). By working at a relatively low SUV concentration, isolated membrane patches could be formed and imaged at 40°C, to confirm the formation of a single stable lipid bilayer in fluid phase. The patches thickness of 4.7±0.1 nm is in line with the expected thickness for such fluid bilayers (81) (Fig. 3D). Increasing the SUV concentration enabled full substrate coverage with a membrane showing only minor defects (Fig. S3).

**Figure 3.**
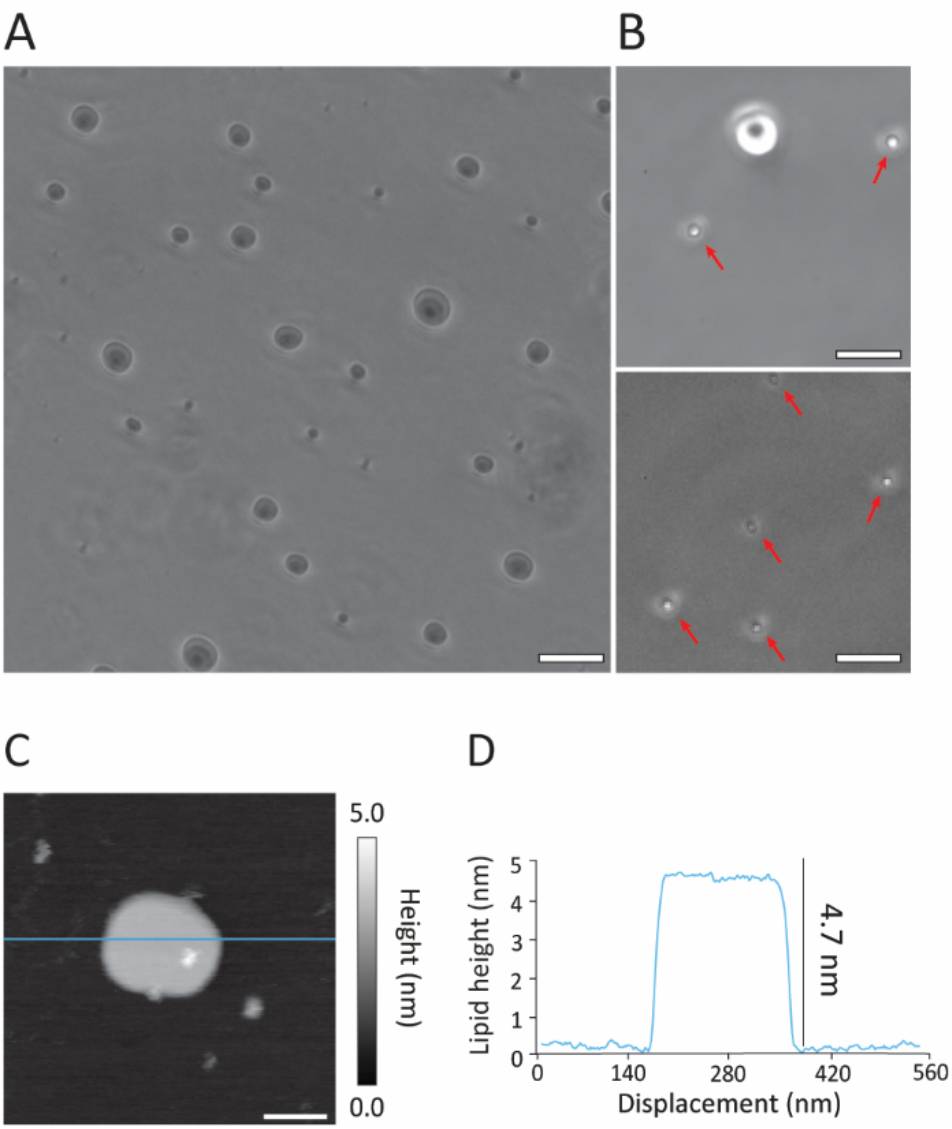
Demonstration of stable membrane formation with candidate mixtures. In bulk solution (A-B), optical microscopy (phase contrast) shows the formation of stable vesicular model membrane systems for the ternary POPE-POPG-CL mixture. Images of spherical ternary mixture’s lipid vesicles were taken with X10 (A) and 40X (B) objectives. 1 μm size vesicles have been indicated with red arrows for clarity (B). The vesicles were stable up to two weeks after the preparation. Stable membranes could also be formed supported on a mica substrate in solution (C-D) with patches of ternary POPE-POPG-CL mixture using low concentrated SUVs solution deposition. AFM imaging (C) reveals the thickness of a typical patch which can be quantified from the associated line profile (D). The scale bars are 50 μm (A - 10X objective), 13 μm (B – 40x objective) and 100 nm (C).

Having confirmed the formation of stable and homogenous SUVs, as well as smooth and stable SLBs we now aim to consider mechanical properties of our model system candidates explicitly. We start by taking into account the Helmholtz free energy of a deformed *E. coli* inner membrane, which consists of the pressure, bending, and stretching/compression elastic energies (82, 83). Ordinarily, *E. coli* cells are under ∼1 atm of osmotic pressure (84, 85), and it is this pressure that leads to mechanical stress in the membrane. While we mentioned such stress can be described via both the bending and stretching energy, because *E. coli* is a spherocylindrical cell 1-3 µm in length and approximately 1 µm in width, the inner membrane curvature is low compared to naturally occurring entropic membrane fluctuations (82, 86), and thus the bending energy can be neglected. We are, therefore, left with the elastic stretching/compression energy of the membrane under mechanical deformation. Fluid membranes cannot support in-plane shear (87) and the associated shear modulus is hence usually taken as zero.

#### Perpendicular compression of the membrane

To probe elastic properties of our candidate model systems under compression we use the nano-indentation of supported membranes with an AFM: the membrane is compressed perpendicularly, squeezed between a nanosphere and a hard substrate. Assuming the membrane to be homogenous and isotropic, its Young’s modulus *Y* (effectively an elasticity modulus in a spring analogy) can be calculated from the out-of-plane compression force at different indentation depths. In practice, it is necessary to take into account the finite thickness of the membrane and the hard substrate, bringing corrections (88, 89) to the established Hertz indentation model for a semi-infinite medium (90). Here we use the following relationship between the applied force *F*_*sphere*_, the indentation depth *δ*, the size of the indenting sphere *R*, and membrane thickness *h* (89):

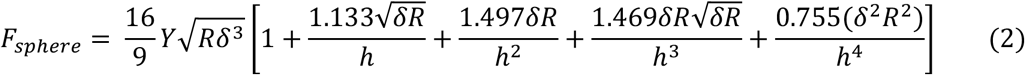

where *δ* ≤ *R*. This formula was derived assuming that the Poisson ratio, which couples in-plane and out-of-plan strain is exactly *ν* = 0.5. In other words, the membrane is assumed incompressible with its volume conserved under compression. This assumption is common for bio-systems (91) and while usually a good approximation, it is not necessarily exact (92, 93). In practice, the indentation of the membrane is carried out with an AFM tip, and while ensuring a linear indentation regime (88) to derive *Y*. We probed the Young’s modulus of mixtures 6A and 9A and compared them with that of the *E. coli* Native and *E. coli Polar* extract obtained in the same manner. The mechanical assays were also performed on two negative controls: a DPPG-based mixture 2B and a pure POPE membrane with both controls being stable when supported by a substrate. In all cases, we distinguished the liquid-disordered (L_d_) and liquid-ordered (L_o_) phases and conducted measurements on both separately. The L_o_ domains are revealed upon cooling of the sampled below its *T*_*m*_, with the L_o_ appearing 0.5 nm to 0.7 nm thicker than the remaining L_d_ phase membrane (Fig. S4A). This is consistent with the expected lipid height variation between the two phases (94, 95).

We then performed so-called force maps (51, 96) whereby force-distance curves – the resistance force experienced by the tip (applied load) as it presses on the membrane – are systematically acquired across randomly selected areas of the membrane (see Fig. S4C-D). From each curve, we immediately get the rupture force *F*_*r*_ (94) necessary for the tip to break through the membrane, and can calculate *Y* from Eq. 3 (Fig. 4). The *E. coli* extracts, and our two candidate mixtures show comparable values on the L_d_ phase (*Y* ∼ 17.5 MPa and *F*_*r*_ *∼* 3.0 nN) and the more ordered L_o_ phase (*Y ∼* 25.8 MPa and *F*_*r*_ ∼ 3.4 nN). All the averages and standard deviations are presented in Table S1. This similarity in mechanical properties is meaningful, as confirmed by the significantly differences derived for the two negative controls: the pure POPE and the DPPG-based samples.

**Figure 4.**
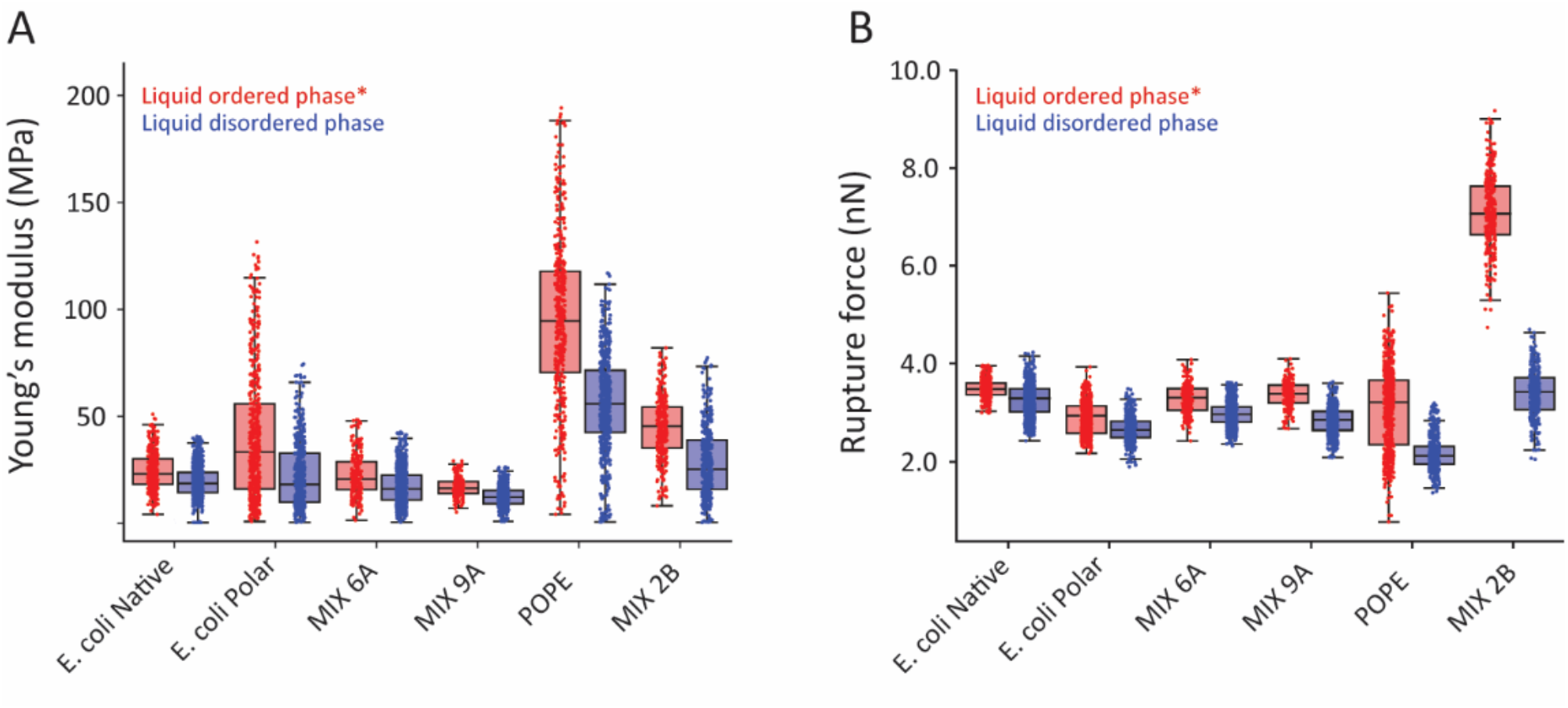
Analysis of the native and candidate membranes using AFM nanomechanical indentation. (A) Membrane average Young modulus, calculated from the indentation region of the curves (elastic indentation) and assuming a spherical tip (radius 12 nm, from manufacturer). The results confirm the mechanical similarities between the E. coli references and two candidate ternary POPE-POPG-CL mixtures. (B) Increasing the indentation force being the elastic region results in the tip puncturing the membrane. The rupture force is simply the force required to break through the lipid bilayer with the AFM tip. For (A) and (B), the data is present as boxplots distinguishing L_o_ and L_d_ phases. In this case, upper and lower whiskers extend to the furthest data point that is within 1.5 times the inter-quartile range, indicating the variability outside the upper and lower quartiles respectively. Note: for membrane composed of a single type of lipid (POPE), the liquid-ordered phase is solid and called ‘gel phase’. The overlayed scatter plot shows the nanomechanical values for each single indentation performed on the membrane. Examples of force maps and measurement principle are illustrated in Fig. S4 B-D.

#### In-plane stretching/compression of the membrane

The AFM measurements use perpendicular indentation under the relatively slow indentation velocities (< 1 µm/s) to provide a single elasticity modulus of the membrane. In many experiments, it is, however, the in-plane stretching/compression elasticity of the membrane that is relevant. Under the same linear elasticity assumptions (82, 97) used previously and assuming the membrane to behave as a 2D material, the in-plane stretching energy is considered (82, 97):

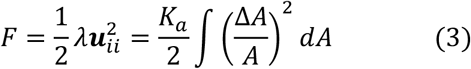

where ***u*** is the strain tensor, λ first Lame coefficient, *K*_*a*_ is equivalent to λ and often referred to as the expansion modulus (98) or elastic area compressibility (99–101) and A is the surface area of the membrane. Since we have assumed the membrane behaves as a homogenous isotropic solid, the simplest model for it, with an equivalent in elastic theory, is a thin plate (98, 102). Then, *K*_*a*_ and *Y* can be related with a well-known relationship:

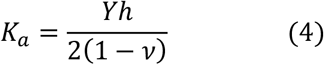

with *h* the membrane thickness and *ν* the Poisson ratio. Taking *ν* = 0.5 and *h* = 4.7 nm (obtained in Fig. 3D) allows calculation of a *K*_*a*_ from the measured *Y*. The result, given in Fig. 5, suggests *K*_*a*_ values slightly lower than the average 0.2 N/m values obtained from previous reports (99–101, 103, 104), where both artificial lipid vesicles and native *E. coli* spheroplast (105) were measured through micropipette aspiration. However, our results still match the correct order of magnitude and fit within the expected range between 0.1 N/m and 0.2 N/m. Thus, both our out-of-plane and in-plane mechanical measurements/estimates support mixtures 6A and 9A as suitable composition to reproduce the biophysical properties of *E. coli*’s inner membrane lipids.

**Figure 5:**
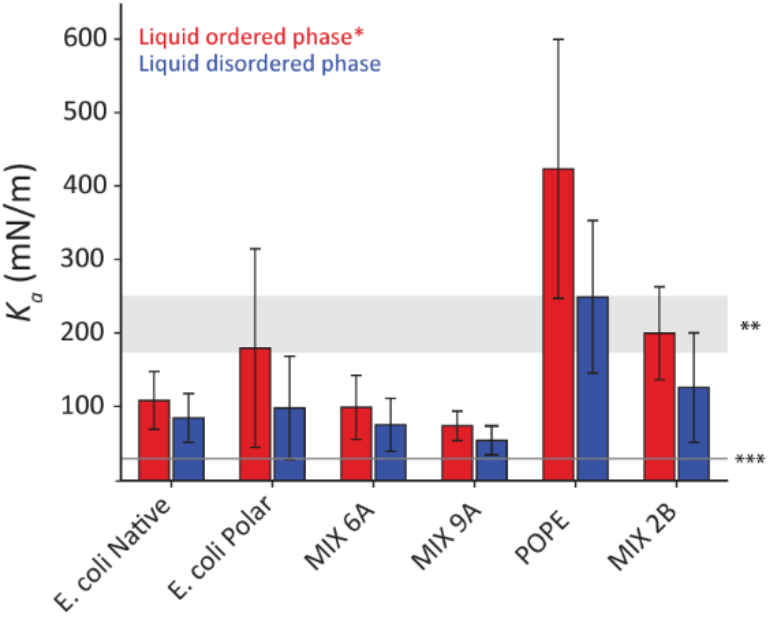
Comparison of the elastic area compressibility, K_a_ values derived from the values obtained for Young’s modulus, Y. The values are derived separately for the Lo and Ld domains of the different candidate membranes using Eq. (4) in conjunction with the data of Fig. 4A and standard deviation has been used as error bar. Values from a previous study (105) have been used to visually compare our K_a_ results. ** K_a_ values for artificial giant lipid vesicles (light grey box), *** K_a_ values for metabolically active E. coli spheroplast (dark grey light), confirming the correct order of magnitude of our results that fits within this range.

## 4 Discussion

In this study we have developed model systems that mimic *E. coli’s* inner membrane lipid composition and mechanical properties based on commercially available lipid mixtures. We find that DPPG is not suitable as a PG source for model *E. coli* membranes under standard growth conditions. Instead, our results indicate the suitability of three ternary mixtures of POPE, POPG and CL as model systems. The mixtures form stable bilayers both in bulk solution and supported. They also match the composition of *E. coli* membrane’s main lipid constituents, the melting temperature of the membrane and its mechanical properties. The fact that the model systems reproduce the main elements of the membranes’ biomechanical proprieties may help future studies where active molecules or forces are at play. For example, we anticipate use of the model systems for investigations of force transduction within membranes, in particular active response to stimuli achieved through integral proteins such as mechanosensitive channels and PIEZO proteins. The transduction and lateral transfer of forces over suitable range and with relevant magnitude, relies on a complex interplay hinging on the local molecular interactions (106) - the local biomechanical properties of the membrane. In the case of *E. coli*, the dynamical shift of *T*_*m*_ as a function of the growth environment indicates a clear correlation between the membrane functionality, the activity of embedded macromolecules and the overall physiological transduction of these stimuli across the cell. Generally, the detection and propagation of mechanical stimuli over controlled distances remains an active research topic with many open questions related to mechano-transduction and the function of these proteins (106, 107). The model systems could also be used to investigate the passive mechanisms behind lipid bilayer asymmetry, an intrinsic property of biological membranes (37), and its effects on the functional features of native membranes. Recent work(108) highlighted how CL can show leaflet preferentiality depending on vesicles curvature, suggesting the possibility of developing model membrane system with controlled compositional asymmetry that could be employed to explore this phenomenon in future work. Additionally, more sophisticated models will be needed to account for the significant local variations in native membrane’s macromolecular content(109). This is important to underpin experimental (110, 111) and computational (107) studies. On the other hand, a simpler bi-component mixture comprising only POPE and POPG could also match native *E. coli* ‘s transition temperature, suggesting its useful for more basic studies where specific molecular interactions or proteins activity is not crucial.

Lastly, we note that the consistency between the L_o_ and L_d_ measurements and the general agreement with existing literature supports our present approach when estimating K_A_, even if it is a simplification, the results need to be taken with caution because the lipid bilayer is not an isotropic 3D material as assumed with the thin plate model.

## Supporting information

Supplementary experiments and controls

## 6 Authors contributions

KV and TP designed the experiment. NT performed all the experiments and carried out the analysis of the results with input from KV and TP. All the authors co-wrote the papers and commented on the results.

## 7 Acknowledgments

Funding from the Engineering and Physical Sciences Research Council SOFI2 Doctoral Training Centre through the EPSRC grant EP/L015536/1 is acknowledged.

