## Supplementary experiments and controls for "A minimalist model lipid system mimicking the biophysical properties of *Escherichia coli’s* membrane"

**Content of the Supplementary Online Material (chronologically):**

- **Figure S1***:* DSC thermographs of MLVs solution to optimize DSC parameters
- **Figure S2***:* DSC thermographs of MLVs solution made of POPG (A) and DPPG (B) based mixtures.
- **Figure S3**. Full coverage and stability of supported lipid bilayers composed of the ternary POPE-POPG-CL mixture (0.3 mg/mL SUVs solution for deposition).
- **Figure S4**. AFM force spectroscopy and force map measurement principles.
- **Table S1.** *Summary* the results obtained from the AFM force maps performed on *the difference* lipid mixtures.
- **References**

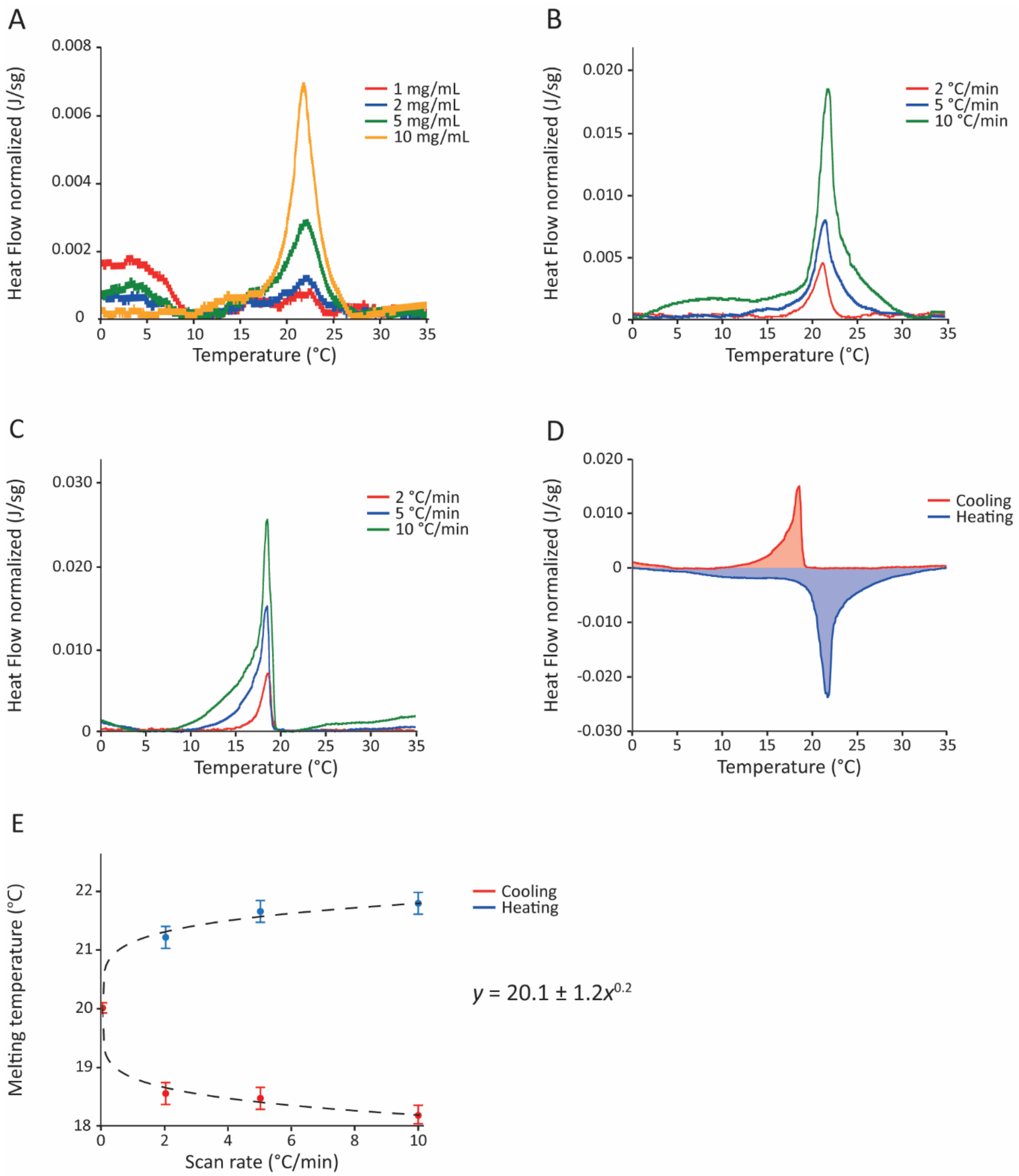


**Figure S2**: DSC thermographs of MLVs solution to optimize DSC parameters. (A) DSC experiments varying MLVs concentration to find the optimal signal to noise (SN) at a scan rate of 5°C/min. (B) DSC experiments varying heating rate to optimise the SN using MLVs solution at 10 mg/mL. Variation of scan rate up to 10°C/min led to small calorimetric differences (melting point variations up to 0.4°C). The same effects are detected on both heating (B) and cooling (C) DSC thermographs, here also acquired using MLVs solution at 10 mg/mL. (D) Direct comparison between cooling and heating DSC experiments with 10 mg/mL solution using a rate of 5°C/min. By using optimal concentration and heating rate, thermograph distortion is limited with relatively small variation of sample's thermodynamic parameters such as the melting point (<3°C), while still maintaining a significant signal to noise ratio. (E) The melting temperature dependence on scan rate, comparing the peaks obtained during heating and cooling can be used to infer the equilibrium transition temperature (rate of zero). The nonlinear dependence of the melting temperature with the scan rate has been previously described (1, 2) as $T_{m,\beta} = T_m + B\beta^z$ where $T_{m,\beta}$ is the measured melting temperature at each scan rate, $T_m$ is the equilibrium or 'true' melting temperature, $\beta$ is the scan rate, and B and z are fitting parameters, B = 1.2 and z = 0.2 after the fit (see inset). Melting temperatures have been reported with standard deviation calculated from 3 different DSC runs per condition.

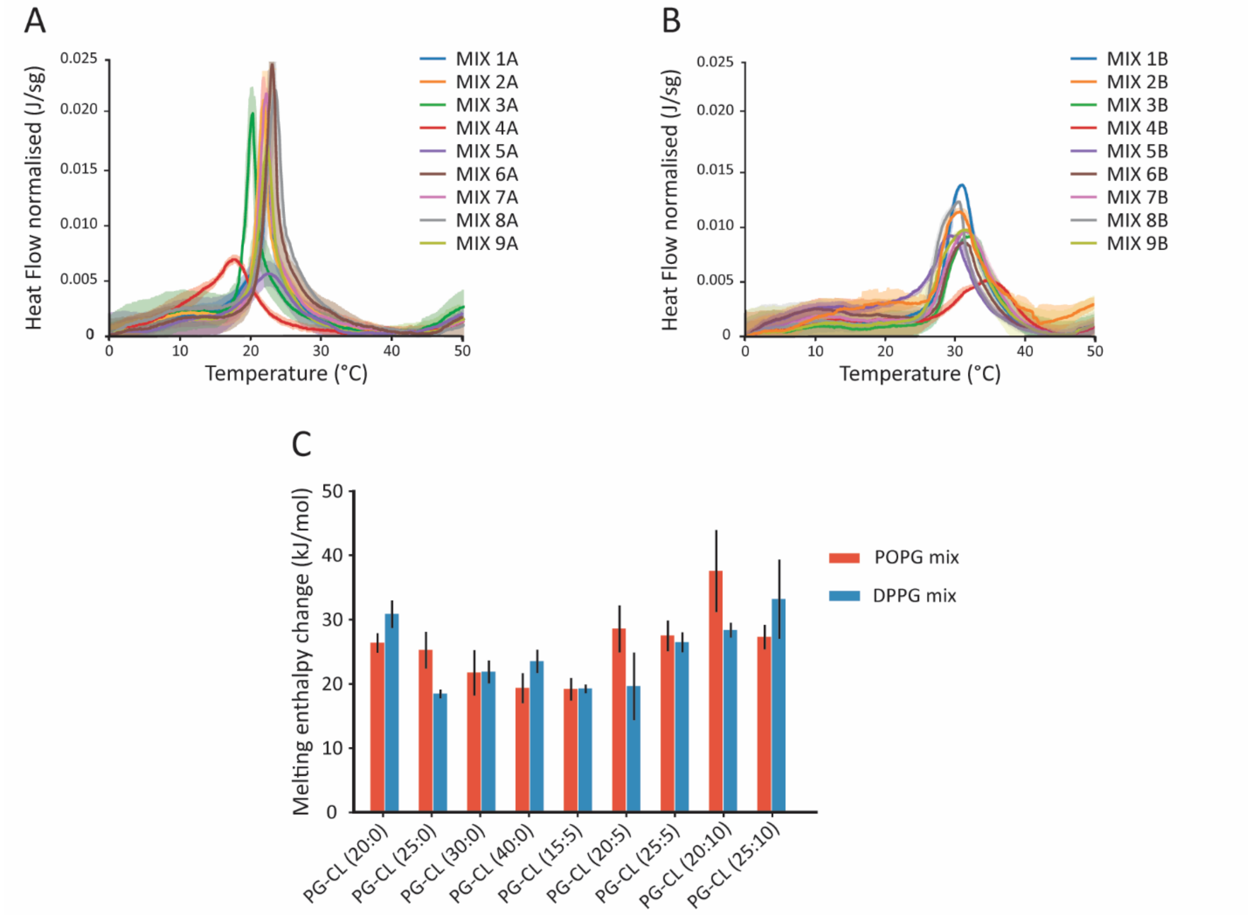


**Figure S2**: DSC thermographs of MLVs solution made of POPG (A) and DPPG (B) based mixtures. The average curve for each mixture has been plotted with standard deviation (shaded areas around average curves). (C) Average melting enthalpy change associated with each DSC main transition peak with standard deviations.

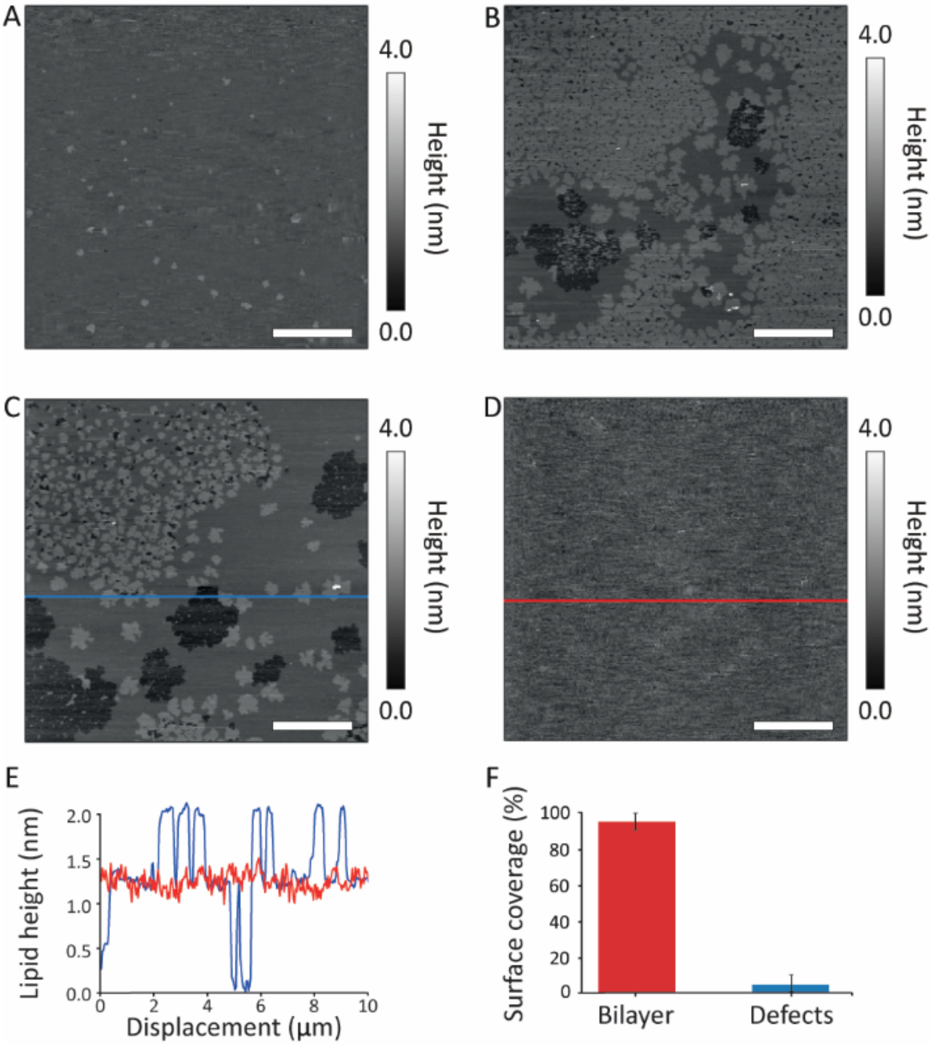


**Figure S3**. Full coverage and stability of supported lipid bilayers composed of the ternary POPE-POPG-CL mixture (0.3 mg/mL SUVs solution for deposition). (A-D) Examples of AFM topographical images of lipid bilayer before (D) and after (A-C) the phase transition. (E) Line profiles showing bilayer and defects in the supported membrane. Blue profile line from Image (C) and red profile line from Image (D). (F) Bar plot representing average bilayer surface coverage calculated over different images (>5) on membranes prepared on different days. The scale bare is 7.5 µm in (A), 5 µm in (B) and 2.5 µm in (C-D).

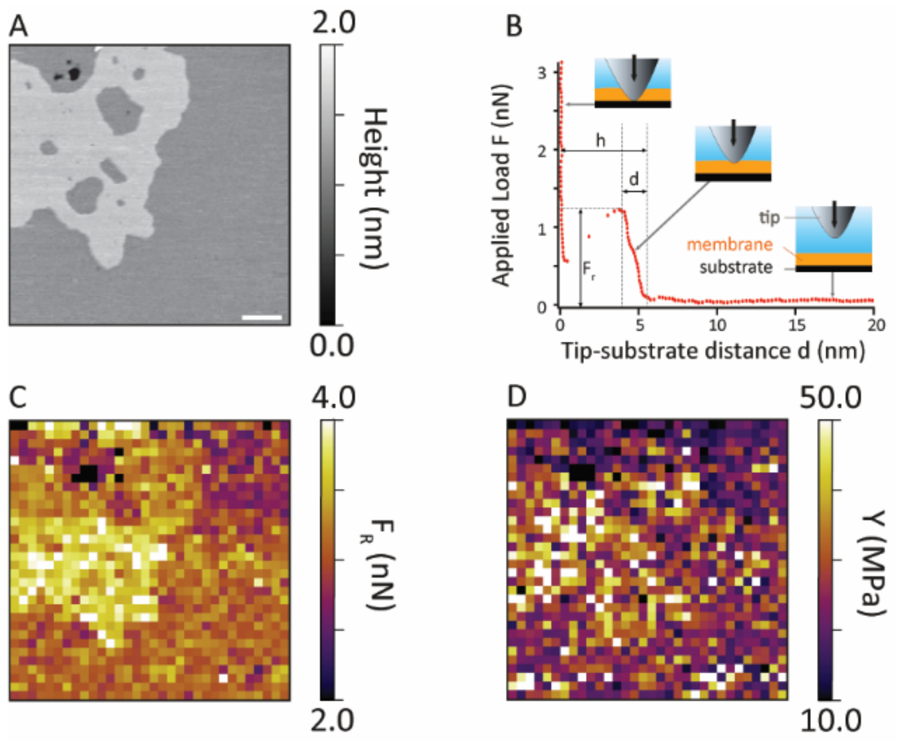


**Figure S4**. AFM force spectroscopy and force map measurement principles. (A) AFM topographical image of the ternary mixture (MIX – 6A) below its melting point. Two lipid phases ($L_o$ and $L_d$ respect. lighter and darker gray) can be seen. (B) Schematic representation of force spectroscopy curve on lipid bilayer with images representing the experimental process. Briefly, when the AFM tip approaches the membrane from the solution, the applied load is zero. Upon contacting the membrane, the applied load begins to increase, compressing and indenting the membrane over a depth d, until the applied load is strong enough to puncture through the bilayer (force $F_R$) and rest on the substrate underneath. From the analysis of the load – tip distance curves, it is possible to estimate the membrane Young's Modulus, *Y,* by fitting the experimental indentation curve with a suitable model (as also discussed in the main text) (3, 4). The total distance *h*~5 nm between the point of membrane contact and the substrate confirms the presence of a single bilayer. (C-D) Force maps performed over the lipid membrane shown in (A) are used to extract $F_R$ and *Y*. The scale bar in (A) is 200 nm.

| Mixture name | Rupture force (nN) | | Young's modulus (MPa) | |
|---|---|---|---|---|
| | Liquid ordered | Liquid disordered | Liquid ordered | Liquid disordered |
| **E. coli Native** | 3.5 ± 0.2 | 3.3 ± 0.3 | 24.3 ± 9.0 | 19.0 ± 7.6 |
| **E. coli Polar** | 2.9 ± 0.3 | 2.7 ± 0.3 | 40.1 ± 30.3 | 22.0 ± 15.9 |
| **MIX – 7A** | 3.3 ± 0.3 | 3.0 ± 0.2 | 22.2 ± 9.9 | 16.9 ± 8.3 |
| **MIX – 9A** | 3.4 ± 0.3 | 2.8 ± 0.3 | 16.6 ± 4.7 | 12.2 ± 4.7 |
| **POPE** | 3.1 ± 0.8 | 2.2 ± 0.3 | 94.3 ± 39.4 | 55.6 ± 23.3 |
| **MIX – 2B** | 7.1 ± 0.8 | 3.4 ± 0.5 | 44.6 ± 14.3 | 28.2 ± 16.8 |

**Table S1.** Summary of the results obtained from the AFM force maps performed on different lipid mixtures. Average and standard deviation are displayed for each mixture and each specific lipid phase (liquid ordered and liquid disordered). Images with an even surface coverage of liquid disordered and liquid ordered phases were chosen. Considering a total of 1024 AFM curves per sample, the values for both the rupture force and the Young's modulus are the average of approximately 500 indentation curves per lipid phase.

4. Hertz, H. 1882. Ueber die Berührung fester elastischer Körper. *Journal für die reine und angewandte. Mathematik*. 92:156–171.
